# Higher seroprevalence of *Entamoeba histolytica* than that of HIV-1 at a voluntary counselling and testing centre in Tokyo

**DOI:** 10.1101/536565

**Authors:** Yasuaki Yanagawa, Mami Nagashima, Hiroyuki Gatanaga, Yoshimi Kikuchi, Shinichi Oka, Keiko Yokoyama, Takayuki Shinkai, Kenji Sadamasu, Koji Watanab

**Author notes:** Corresponding author information: Koji Watanabe, AIDS Clinical Center, National Center for Global Health and Medicine. 1-21-1, Toyama, Shinjuku-ku, Tokyo 162-8655, Japan. These authors contributed equally to this article (Yasuaki Yanagawa, Mami Nagashima). Email addresses: Yasuaki Yanagawa, Mami Nagashima, Hiroyuki Gatanaga, Yoshimi Kikuchi, Shinichi Oka, Keiko Yokoyama, Takayuki Shinkai, Kenji Sadamasu, Koji Watanabe.

## Abstract

**Background:** Amebiasis, which is caused by *Entamoeba histolytica*, is a re-emerging public health issue owing to sexually transmitted infection (STI) in Japan. However, epidemiological data are quite limited.

**Methodology:** To reveal the relative prevalence of sexually transmitted *E. histolytica* infection to other STIs, we conducted a cross-sectional study at a voluntary counselling and testing (VCT) centre in Tokyo. Seroprevalence of *E. histolytica* was assessed according to positivity with an enzyme-linked immunosorbent assay for *E. histolytica*-specific IgG in serum samples collected from anonymous VCT clients.

**Principal Findings:** Among 2,083 samples, seropositivity for *E. histolytica* was 2.64%, which was higher than that for HIV-1 (0.34%, p < 0.001) and comparable to that for syphilis (rapid plasma reagin (RPR) 2.11%, p = 0.31). Positivity for *Chlamydia trachomatis* in urine by transcription-mediated amplification (TMA) was 4.59%. Seropositivity for *E. histolytica* was high among RPR-or *Treponema pallidum* hemagglutination (TPHA)-positive individuals and it was not different between clients with and without other STIs. Both seropositivity of *E. histolytica* and RPR were high among male clients. The seropositive rate for anti-*E. histolytica* antibody was positively correlated with age. TMA positivity for urine *C. trachomatis* was high among female clients and negatively correlated with age. Regression analysis identified that male sex, older age, and TPHA-positive results are independent risk factors of *E. histolytica* seropositivity.

**Conclusions:** Seroprevalence of *E. histolytica* was 7.9 times higher than that of HIV-1 at a VCT centre in Tokyo, with a tendency to be higher among people at risk for syphilis infection.

**Author summary:** Amebiasis caused by *Entamoeba histolytica* is an increasingly prevalent sexually transmitted infection (STI) in Japan; however, relative to other STIs, the prevalence of *E. histolytica* has not been fully assessed. We investigated the seropositivity of *E. histolytica* using serum samples from 2,083 clients of a voluntary counselling and testing centre in Tokyo. *E. histolytica* seroprevalence (2.64%) was 7.9 times higher than that of HIV-1 (0.31%) and the same as that of syphilis (rapid plasma reagin: 2.11%). Logistic regression analysis showed that *E. histolytica* seroprevalence tended to be higher among individuals who were male, older, and positive in *Treponema pallidum* hemagglutination. These results strongly suggest that public health interventions should be considered to control sexual transmission of *E. histolytica* infection, which is currently neglected in Japan.

## INTRODUCTION

Amebiasis is an enteric protozoa infection caused by *Entamoeba histolytica*. Up to 80% of *E. histolytica* infections are asymptomatic but persistent; the remainder result in the development of invasive diseases, such as colitis and liver abscess [1]. Asymptomatically infected individuals represent a risk to the community because they are a source of new infections. Transmission occurs via the oral–faecal route. It has long been believed that amebiasis is only endemic in developing countries where food and water are frequently contaminated with human faeces, or that it occurs among travellers to or immigrants from these countries [1, 2]. However, in the previous two decades, it has been reported that cases of amebiasis have been rapidly increasing and have become a re-emerging infectious disease in developed countries of East Asia and in Australia [3-8]. Human-to-human transmission occurs via direct sexual contact, such as oral– anal sexual contact and contact among men who have sex with men in these countries [9, 10]. Under such circumstances, it is essential to identify individuals who are asymptomatic but chronically infected with *E. histolytica* and who thus represent sources of new infection, for the epidemiologic control of sexually transmitted *E. histolytica* infection. However, little epidemiological data is currently available in Japan, other than that from National Epidemiological Surveillance of Infectious Diseases (NESID), which only reports clinically diagnosed “symptomatic” cases. Moreover, it is critical to understand the epidemiology of sexually transmitted *E. histolytica* infection before the upcoming Tokyo Olympics in 2020, which could serve as a source of the rapid spread of such neglected communicable diseases.

In the present study, we investigated the seroprevalence of *E. histolytica* at a voluntary counselling and testing (VCT) centre in Tokyo, in comparison with the prevalence of other sexually transmitted infections (STIs). In addition, we discuss future strategies for the epidemiologic control of sexually transmitted *E. histolytica* infection.

## METHODS

### Setting

Tokyo, the capital city of Japan, is located on the Pacific on the eastern coast of Honshu, the largest of the four main islands comprising Japan. According to the national surveillance system, the annual number of HIV tests performed and the incidence rates of HIV infection are higher in Tokyo than those of other prefectures [11]. The Tokyo Metropolitan Minami Shinjuku Testing – Counselling Centre is the largest HIV testing centre in Tokyo, and it is very close to a town in Shinjuku with a large population of men who have sex with men (MSM) [12]. Because there are more MSM who visit this centre to undergo testing for HIV and other STIs, the incidence rate of HIV infection at this centre is higher than that of other public health centres in Tokyo [13].

### Study population, samples, and ethics issues

The design of this study was a cross-sectional study. The total 2,083 serum samples used in this study were collected at the Tokyo Metropolitan Minami Shinjuku Testing – Counselling Centre where more than 10,000 anonymous clients seek HIV-1 screening tests each year. Collected samples are transferred to the Tokyo Metropolitan Institute of Public Health for laboratory testing, then stored at 4°C. Fourth generation HIV-1 screening is performed routinely throughout the year. However, in 2 months of the year (e.g., June and December in the case of 2017), the Tokyo Metropolitan Government intensifies STI screening, and rapid plasma reagin (RPR) and *Treponema pallidum* hemagglutination (TPHA) tests for syphilis screening are additionally performed for all clients. In addition, urinary sampling and transcription-mediated amplification (TMA) assay testing for *Chlamydia trachomatis* and *Neisseria gonorrhoeae* are performed for clients who are willing to undergo these tests. Therefore, we assessed the seroprevalence of anti-*E. histolytica* antibody using stored serum samples collected in June and December of 2017 and compared this with the positivity for other STIs in the present study. In the present study, there was no selection bias or missing data.

This study was approved by the ethics committee of the National Center for Global Health and Medicine (NCGM-2302) and Tokyo Metropolitan Institute of Public Health (29-875). All protocols for this study were conducted in accordance with the Declaration of Helsinki.

### Laboratory testing

The presence of anti-*E. histolytica* antibody was detected using a commercially available ELISA kit (*Entamoeba histolytica* IgG-ELISA; GenWay Biotech, Inc., San Diego, CA. USA). All procedures were performed according to the manufacturer’s instructions. In brief, diluted serum samples (100X dilution in IgG sample diluent) as well as 5 control samples, consisting of 1 substrate blank, 1 negative control, 2 cut-off controls, and 1 positive control, were applied to 96-well plates pre-treated with *E. histolytica* antigen and incubated at 37°C for 1 hour. After washing the plates using washing solution, 100 μL of *E. histolytica* Protein A conjugate was added to all wells except the substrate blank and incubated for 30 minutes in the dark. After a second wash, TMB (3,3’,5,5’-Tetramethylbenzidine) substrate solution was added to all wells. After a 15-minute incubation, 100 μL of stop solution was applied to the plates, and absorbance of the specimen was then read at 450/620 nm using a spectrometer.

### Statistical analysis

Of the total samples tested in each STI screening test, the proportion of seropositive blood and urine samples are presented with 95% confidence interval (CIs) calculated using the Wilson– Brown method. The seroprevalence of *E. histolytica* was compared with that of other sexually transmitted infections using Fisher’s exact test. To determine the trend of seropositivity among age groups, we used the chi-square test for trend. Statistical significance was defined as a two-sided p value < 0.05. All statistical analyses were conducted using GraphPad Prism (GraphPad Software, La Jolla, CA, USA). Logistic regression analysis for identification of factors influencing *E. histolytica* seropositivity was performed using Stata (StataCorp LLC., College Station, TX, USA).

## RESULTS

### Study population and seroprevalence of *E. histolytica* at a voluntary counselling and testing centre in Tokyo

In total, 2,083 samples were analysed. The average age of clients was 35.2 (95% CI: 34.8– 35.7) years, and 70.8% (1474/2083) were male (Fig 1). The overall seropositivity for *E. histolytica* was 2.64%; this was significantly higher than that for HIV-1 (0.34%) and the comparable level as that for syphilis by RPR (2.11%) (Fig 2A). The positive rate of urinary TMA for *C. trachomatis* (4.59%) was higher than the seropositivity for *E. histolytica*; however, urinary TMA testing for *C. trachomatis* and *N. gonorrhoeae* was only carried out in 69.0% (1,437/2,083) of clients, i.e., those who were willing to undergo TMA testing. These results suggest that *E. histolytica* is a more common STI than HIV-1 in Tokyo and is at a level comparable to that of syphilis infection. Interestingly, all individuals who were seropositive for *E. histolytica* were seronegative for HIV-1 (Fig 2B). Furthermore, the seropositive rate for *E. histolytica* was significantly higher among people who were seropositive for syphilis infection (by both RPR and TPHA) than among those who were seronegative for syphilis; no significant differences in *E. histolytica* seropositivity were seen according to TMA positivity for *C. trachomatis*. These results indicate that *E. histolytica* infection is spreading among people at risk for syphilis infection.

**Figure 1.**
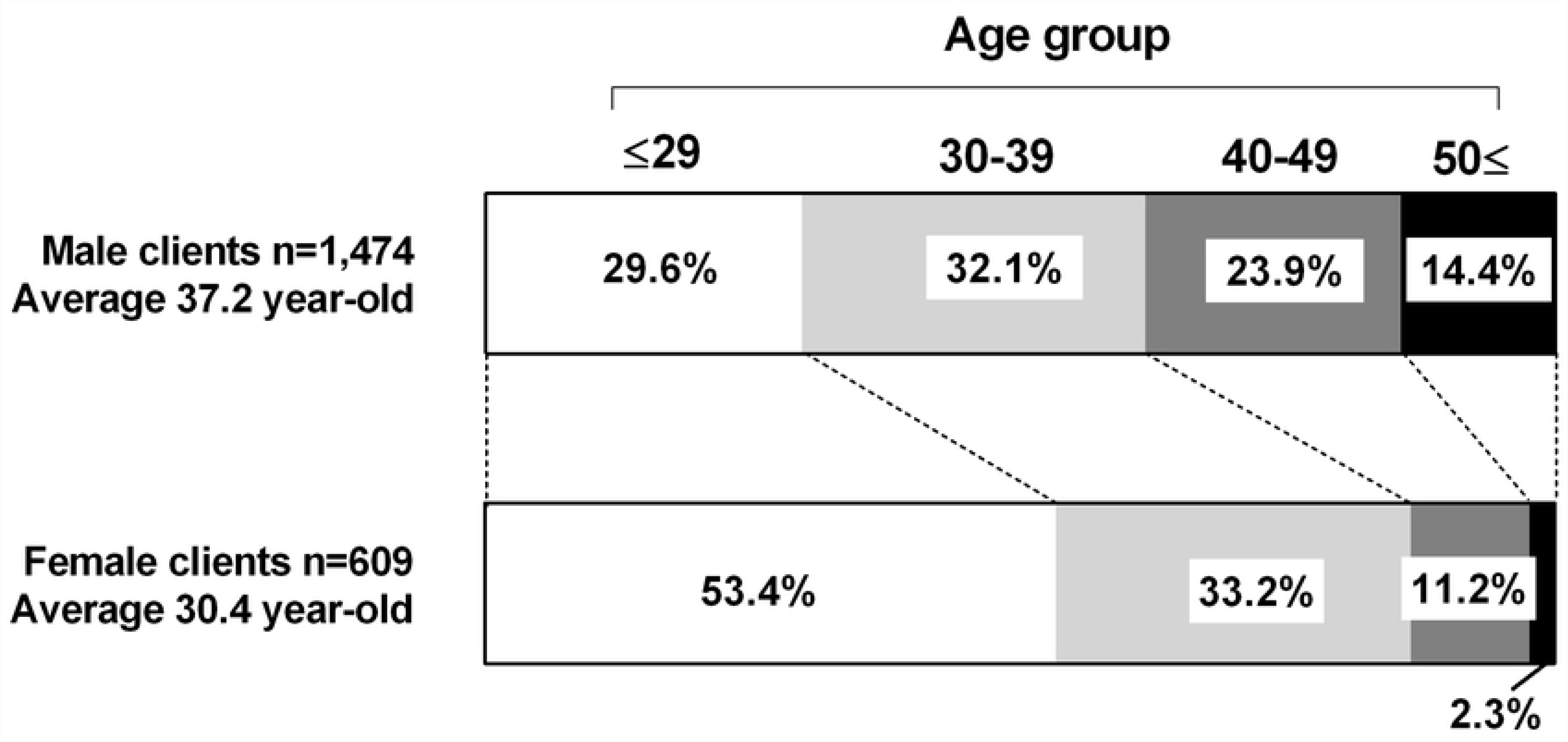
Proportion of clients in each age group among men and women. The average age among female clients was significantly lower than that in male clients (p < 0.001). The proportion of clients aged 29 years or less among female clients was 53.4% whereas that in male clients was only 29.6%.

**Figure 2.**
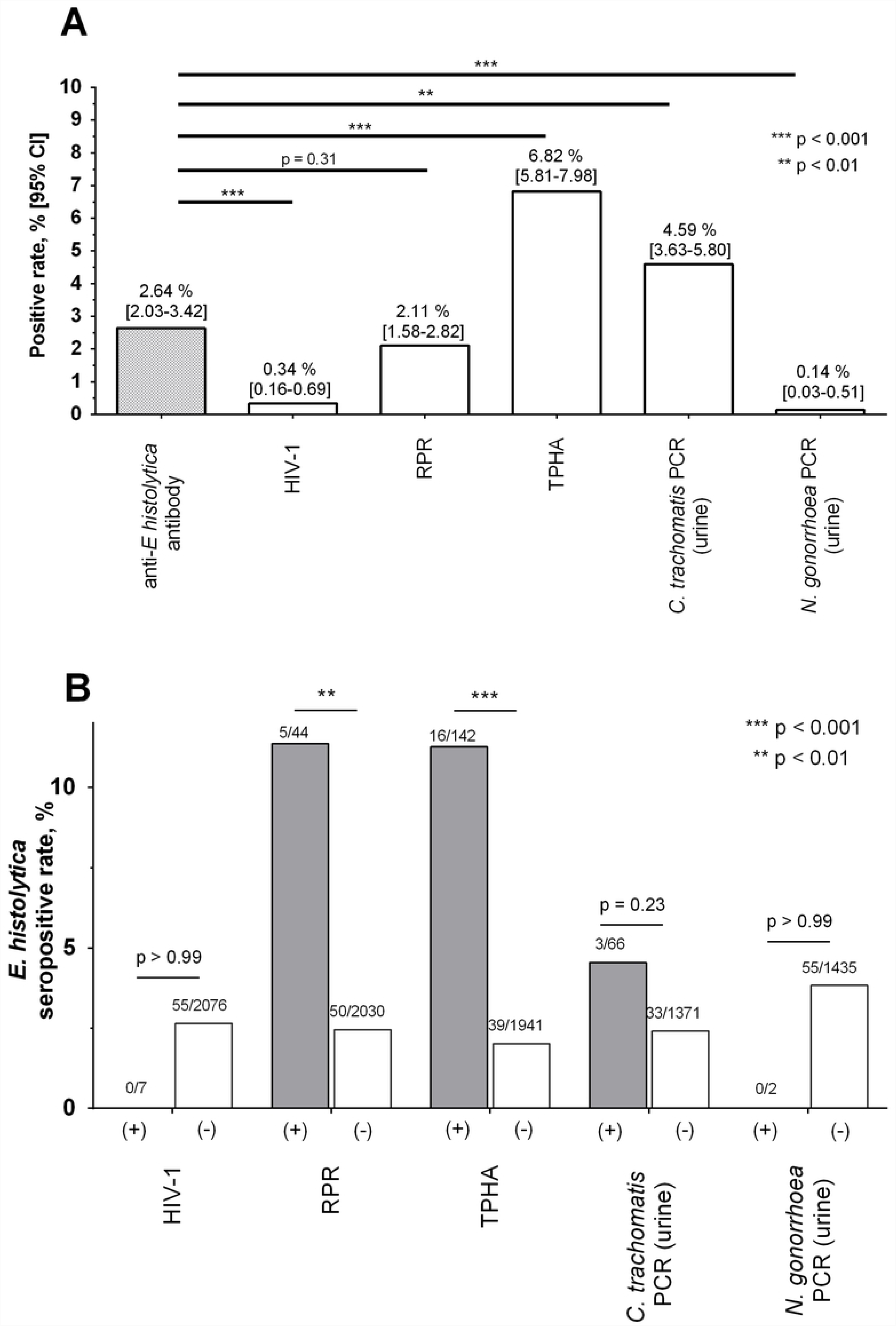
Seropositivity for *Entamoeba histolytica* and other sexually transmitted infections (STIs) in Tokyo. Serologic testing results (anti-*E. histolytica* antibody, HIV-1, RPR, and TPHA) were obtained for 2,083 clients of a voluntary counselling and testing centre in June and December of 2017. Results of urinary TMA for *Chlamydia trachomatis* and *Neisseria gonorrhoeae* were available for 1,437 clients who agreed to testing. All statistics were calculated using Fisher’s exact test. (A) The seropositive rate for *E. histolytica* was compared with those of other STIs. (B) Comparison of seropositivity for *E. histolytica*, with and without other STIs. Abbreviations: CI, confidence interval; RPR, rapid plasma reagin; TPHA, *Treponema pallidum* hemagglutination; TMA, transcription-mediated amplification.

### Differences in seropositivity by sex and age group

Next, we compared positivity for STIs between male and female clients. Seropositivity for *E. histolytica* was significantly higher in male (3.46%) than in female (0.66%) clients, as seen for syphilis infection (RPR: 2.78% vs. 0.49% and TPHA: 9.29% vs. 0.82%) (Fig 3A). The proportion of urinary TMA results positive for *C. trachomatis* was significantly higher in female (8.77%) than male (2.65%) clients. However, it is difficult to simply compare the TMA positivity by sex because persistent, asymptomatic *C. trachomatis* infection of the urinary tract occurs more frequently in females [14-17]. Moreover, the age of female clients was significantly lower than that of males, and the proportion of clients aged 29 years or less in females was 53.4% whereas that in males was only 29.6% (Fig 1). These results indicate that both male and female clients in this study are at risk for STIs; however, the predominant pathogens might differ between relatively older males (*E. histolytica* and *T. pallidum*) and relatively younger females (*C. trachomatis*).

**Figure 3.**
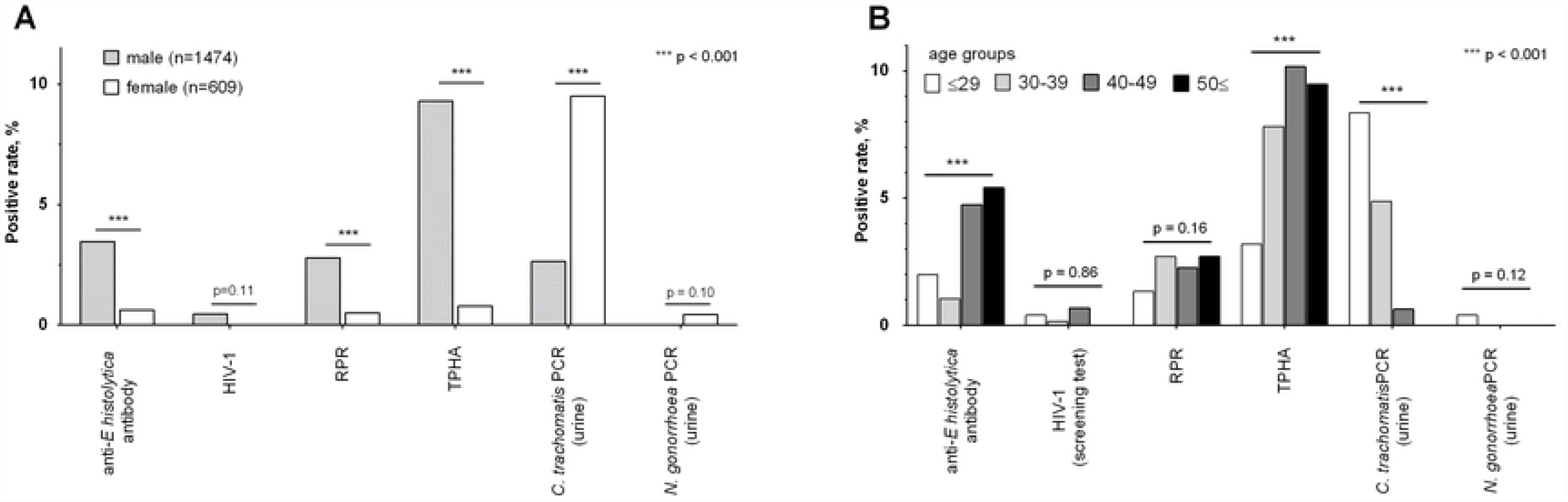
Positive rate of sexually transmitted infections (STIs) by sex and age group. (A) Positive rate of *Entamoeba histolytica* and other STIs were compared between male (n = 1474) and female (n = 609) clients using Fisher’s exact test. (B) Seropositivity for *E. histolytica* and RPR, and TMA positivity for *Chlamydia trachomatis* were calculated for clients of different age groups (serum, urine samples): 29 years or younger (752, 503), 30–39 years (666, 453), 40–49 years (443, 315), and 50s or older (222, 167). Correlation between age and positivity was calculated using the chi-square test for trend. Abbreviations: RPR, rapid plasma reagin test; TPHA, treponema pallidum hemagglutination; TMA, transcription-mediated amplification.

To determine the trend of *E. histolytica* seropositivity by age, we compared seropositivity for *E. histolytica* in different age groups. Interestingly, seropositivity for anti-*E. histolytica* antibody and RPR was highest among clients aged 50 years or older (5.41% and 2.70%, respectively). Moreover, a positive correlation was observed between age and seropositivity for *E. histolytica* (Fig 3B). Positive urinary TMA for *C. trachomatis* was highest among clients aged 29 years or younger (8.35%) and showed a negative correlation with age. These results are consistent with national surveillance data, in which diagnosed cases of *Chlamydia* infection have a peak in the 20s [18], whereas the median age of reported cases of amebiasis is relatively high (50 years in men and 40 years in women) [5, 16]. Considering these findings, *E. histolytica* infection might be more prevalent among relatively older age groups (40 years or more) whereas *Chlamydia* infection is more prevalent in relatively younger populations.

### Risk of seropositivity for *E. histolytica*

Finally, to identify the risk factors of seropositivity for *E. histolytica*, we performed logistic regression analysis using data of client characteristics and the results of STI screening tests. Univariate and multivariate regression analyses revealed that male sex, a history of syphilis infection (by TPHA), and older age were independent risk factors of seropositivity for *E. histolytica* (Table 1). In particular, age 40 years or older was a high-risk factor of seropositivity for *E. histolytica* (odds ratio 3.31 in people aged less than 40 years, p value < 0.001 by univariate analysis; data not shown). In addition, univariate analysis showed that positive RPR was a high-risk factor for *E. histolytica* seropositivity; however, this was diminished in multivariate analysis owing to the strong association with TPHA positivity. Univariate analysis using preliminary urinary TMA data of 1,437 participants showed that positivity for *C. trachomatis* in the urine had no impact on *E. histolytica* seropositivity (Table 1). We could not include HIV-1 serology and TMA positivity for *N. gonorrhoeae* in urine in the logistic regression analyses because no clients who were HIV-1 seropositive or positive for *N. gonorrhoeae* by TMA were also seropositive for *E. histolytica*.

**Table 1.** Impact of individual characteristics on seropositivity for *Entamoeba histolytica*, Tokyo.

|  |  | Univariate analysis |  | Multivariate analysis*** |  |
| --- | --- | --- | --- | --- | --- |
|  |  | OR<br>(95% CI) | p value | OR<br>(95% CI) | p value |
| Sex (Male) |  | 5.42<br>(1.95–15.06) | < 0.001 | 3.17<br>(1.10–9.07) | 0.032 |
| Older age<br>(by 10-year age groups) |  | 1.66<br>(1.33–2.08) | < 0.001 | 1.49<br>(1.17–1.90) | 0.001 |
| HIV-1 positive |  | ND** |  |  |  |
| Syphilis<br>infection | RPR positive | 5.1<br>(1.93–13.49) | 0.006 | 1.26<br>(0.41–3.89) | 0.693 |
|  | TPHA positive | 6.19<br>(3.37–11.39) | < 0.001 | 4.30<br>(2.11–8.76) | < 0.001 |
| Urine *C. trachomatis* (TMA)<br>positive* |  | 1.93<br>(0.58–6.47) | 0.326 |  |  |
| Urine *N. gonorrhoeae* (TMA)<br>positive* |  | ND** |  |  |  |

\* Data of urinary TMA testing available only for 69.0% (1,437 of 2,083) of total clients.
\*\* Odds ratios could not be determined in logistic regression analysis because all clients who were HIV-1 positive and/or positive for gonorrhoea by TMA were *E. histolytica* seronegative.
\*\*\* Multivariate analysis for age and sex, plus factors with $p < 0.05$ in univariate analysis.

Abbreviations: OR, odds ratio; RPR, rapid plasma reagin; TPHA, *Treponema pallidum* hemagglutination; TMA, transcription-mediated amplification; ND, not determined.

## DISCUSSION

The most important finding of the present study was that the seroprevalence of *E. histolytica* was significantly (7.9 times) higher than that of HIV-1 and it was comparable to that for syphilis (by RPR). Certainly, it is difficult to simply compare seropositivity among these three tests; the HIV-1 screening test continues to be positive for a person’s entire life whereas positivity in RPR and anti-*E. histolytica* antibody tests indicate current or recent infection [17, 18]. However, these results strongly indicate that the endemicity of *E. histolytica* in Tokyo is higher than that of HIV-1 and close to the level of syphilis. In contrast to our seroprevalence data, the national surveillance data of Japan from NESID pragmatically show that the annual number of diagnosed cases of amebiasis (1,151 in 2016) is not only much lower than that of syphilis (4,575 cases), it is also lower than that of HIV-1 (1,443 cases) [18, 19]. Our results suggest that the endemicity of amebiasis in Japan is currently underestimated, thereby remaining a neglected disease in Japan despite frequently reported life-threatening cases of amebiasis [22-25]. Interestingly, in the present study, all individuals who were seropositive for *E. histolytica* were HIV-1 negative whereas regression analysis identified that seropositivity for syphilis by TPHA was an independent risk factor of a positive result for anti-*E. histolytica* antibody. Previous reports have emphasized the high seroprevalence of *E. histolytica* [26] and increasing number of amebiasis cases [27-29] among individuals with HIV-1 infection. Although the epidemiological trend of *E. histolytica* among HIV-1-positive individuals could not be assessed in this study owing to the small number of clients who were positive for HIV-1, it should be noted that sexually transmitted *E. histolytica* infection is currently spreading even among HIV-1-negative populations, as we indicated in our previous hospital-based cross-sectional analysis [30]. Currently, screening for *E. histolytica* is not routinely performed at VCT centres in Japan; however, public health interventions should be considered to control sexually transmitted *E. histolytica* infection.

The clinical significance of seropositivity for *E. histolytica* remains unclear and is beyond the scope of this paper. Serologic testing is a sensitive diagnostic method for symptomatic invasive amebiasis; however, positive results are also obtained for recent infections, up to the previous several years [23]. However, we previously reported that 70.4% of *E. histolytica*-seropositive individuals did not have any amebiasis-related symptoms nor any history of treatment for amebiasis. Interestingly, 20% of such individuals in a Japanese HIV-1 cohort developed symptomatic invasive amebiasis within a 1-year follow-up period [23]. In another cross-sectional analysis, we also reported that ulcerative lesions owing to *E. histolytica* in the large intestine are frequently identified (7/18, 38.9%) by colonoscopy among asymptomatic individuals who are *E. histolytica* seropositive whereas these rarely (1/53, 1.9%) occur among *E. histolytica*-seronegative people [31]. Serologic screening for *E. histolytica* at VCT centres, followed by diagnosis of subclinical *E. histolytica* infection by colonoscopy and treatment at a referral hospital, is one possible public health strategy against sexually transmitted *E. histolytica* infection. However, we must assess the utility of serologic testing for the screening of asymptomatic *E. histolytica* in well-designed prospective analyses in the future.

The present study has some limitations that should be considered. First, this preliminary investigation was a cross-sectional study of anonymous clients at a VCT centre. We could not assess risk behaviour or sexual behaviour with respect to seropositivity for *E. histolytica* owing to a lack of detailed data on the characteristics of clients. In addition, the study periods were 2 months apart owing to the availability of data for not only HIV-1 but also other STIs (serum tests for syphilis and urine tests for chlamydia and gonorrhoea). We could not exclude the possibility of selection bias of clients, such as those who undergo repeat testing. In future, intensifying STI screening services at VCT centres should be considered, for epidemiological control of these prevalent STIs in Japan. Second, anti-*E. histolytica* antibody was screened using stored serum. Long periods of storage could lead to lower sensitivity of serologic tests, resulting in underestimation of the seroprevalence of *E. histolytica*. Third, we obtained a considerably lower seropositive rate for *E. histolytica* among female clients (0.66%, 4/609) than that among males (3.46%, 51/1,474). This probably results from the fact that VCT centres may not be appropriate for identifying female populations at high risk for *E. histolytica* infection; our female clients were relatively younger and had lower seropositivity in RPR and TPHA tests. More appropriate sampling locations should be identified, such as STI clinics that are visited by female commercial sex workers [32].

In conclusion, among clients of a VCT centre in Tokyo, seropositivity for *E. histolytica* was 7.9 times higher than that of HIV-1 and tended to be high among individuals at risk of syphilis infection. Active detection and treatment of asymptomatic cases of *E. histolytica* infection should be considered for the epidemiologic control of sexually transmitted *E. histolytica* infection in Japan.

## Acknowledgements

This work was supported by the Emerging/Re-emerging Infectious Diseases Project of Japan from the Japan Agency for Medical Research and Development (AMED), and a grant from the National Center for Global Health and Medicine (29-2013). We thank Analisa Avila, ELS, of Edanz Group (www.edanzediting.com/ac) for editing a draft of this manuscript.

## Supporting Information

**S1 Fig. Figure 1. Proportion of clients in each age group among men and women.** The average age among female clients was significantly lower than that in male clients (p < 0.001). The proportion of clients aged 29 years or less among female clients was 53.4% whereas that in male clients was only 29.6%.

**S2 Fig. Figure 2. Seropositivity for *Entamoeba histolytica* and other sexually transmitted infections (STIs) in Tokyo.** Serologic testing results (anti-*E. histolytica* antibody, HIV-1, RPR, and TPHA) were obtained for 2,083 clients of a voluntary counselling and testing centre in June and December of 2017. Results of urinary TMA for *Chlamydia trachomatis* and *Neisseria gonorrhoeae* were available for 1,437 clients who agreed to testing. All statistics were calculated using Fisher’s exact test. (A) The seropositive rate for *E. histolytica* was compared with those of other STIs. (B) Comparison of seropositivity for *E. histolytica*, with and without other STIs. Abbreviations: CI, confidence interval; RPR, rapid plasma reagin; TPHA, *Treponema pallidum* hemagglutination; TMA, transcription-mediated amplification.

**S3 Fig. Figure 3. Positive rate of sexually transmitted infections (STIs) by sex and age group.** (A) Positive rate of *Entamoeba histolytica* and other STIs were compared between male (n = 1474) and female (n = 609) clients using Fisher’s exact test. (B) Seropositivity for *E. histolytica* and RPR, and TMA positivity for *Chlamydia trachomatis* were calculated for clients of different age groups (serum, urine samples): 29 years or younger (752, 503), 30–39 years (666, 453), 40–49 years (443, 315), and 50s or older (222, 167). Correlation between age and positivity was calculated using the chi-square test for trend. Abbreviations: RPR, rapid plasma reagin test; TPHA, treponema pallidum hemagglutination; TMA, transcription-mediated amplification.

**S1 Checklist**: STROBE Checklist for cross-sectional studies

